# A genome-wide enrichment screen identifies NUMA1-loss as a resistance mechanism against mitotic cell-death induced by BMI1 inhibition

**DOI:** 10.1101/2019.12.24.887851

**Authors:** Santiago Gisler, Ana Rita R. Maia, Gayathri Chandrasekaran, Maarten van Lohuizen

**Affiliations:** Division of Molecular Genetics, Oncode and The Netherlands Cancer Institute, Amsterdam, The Netherlands; Division of Cell Biology, The Netherlands Cancer Institute, Amsterdam, The Netherlands

## Abstract

BMI1 is a core protein of the polycomb repressive complex 1 (PRC1) that is overexpressed in several cancer types, making it a promising target for cancer therapies. However, the underlying mechanisms and interactions associated with BMI1-induced tumorigenesis are often context-dependent and complex. Here, we performed a drug resistance screen on mutagenized human haploid HAP1 cells treated with the BMI1 inhibitor PTC-318 to find new genetic and mechanistic features associated with BMI1-dependent cancer cell proliferation. Our screen identified NUMA1-mutations as the most significant inducer of PTC-318 cell death resistance. Independent validations on NUMA1-proficient HAP1 and non-small cell lung cancer cell lines exposed to BMI1 inhibition by PTC-318 or *BMI1* knockdown resulted in cell death following mitotic arrest. Interestingly, cells with CRISPR-Cas9 derived *NUMA1* knockout also showed a mitotic arrest phenotype following BMI1 inhibition but, contrary to cells with wildtype NUMA1, these cells were resistant to BMI1-dependent cell death. The current study brings new insights to BMI1 inhibition-induced mitotic lethality in cancer cells and presents a previously unknown role for NUMA1 in this process.

## Introduction

The chromatin-modifying Polycomb-group proteins are critical epigenetic transcriptional repressors controlling cell fate decisions, such as self-renewal and differentiation of stem cells, as well as tumorigenesis, primarily through the repression of downstream genes (1–3). B lymphoma Mo-MLV insertion region 1 homolog (BMI1), an essential protein of the polycomb repressive complex 1 (PRC1), was first identified as an oncogene, inducing lymphomas in mice by co-operating with c-MYC (4,5). The protein is often expressed in stem cells, and several reports have implicated its overexpression in cancer stem cell maintenance and the progression of different types of cancers (6–8). By contrast, regulation of BMI1 using inhibitors or short hairpin RNAs results in cellular senescence or apoptosis of several types of cancer cells (9–13), and sensitizes tumor cells to cytotoxic agents or radiation (14,15). Because of this, BMI1 is an attractive target for future clinical therapies of different cancers.

BMI1 overexpression is a well-established inducer of cancer cell proliferation and resistance to cancer drug treatments of various cancer cell lines (16–18), highlighting the potential of specific BMI1 inhibitors. However, although BMI1 inhibition results in growth reduction and cell death of different cancer cell lines, the underlying mechanisms are often context-dependent and uncertain (11,13,19). As a result, little is known about the genetic interactions and variations involved in BMI1 inhibition-derived lethality or the subsequent resistance.

In the present study, we performed a genome-wide screen for gene disruptions that could result in resistance to pharmacological inhibition of BMI1 by exposing mutagenized human haploid HAP1 cells to low concentrations of the BMI1-inhibitor AB057609107 (PTC-318). PTC-318 is a new inhibitor of BMI1, developed by PTC Therapeutics, USA, that is designed to regulate BMI1 expression post-transcriptionally. We show that reduction of BMI1 levels by short hairpin RNA (shRNA) or inhibition with the small molecule inhibitor PTC-318 significantly reduced cell viability of different cancer cell lines. Through our genetic screen, we unveiled new genes that rescue cell death induced by BMI1 inhibitors and selected NUMA1 for follow-up studies. We identified a novel mitotic mechanism underlying BMI1-associated lethality using CRISPR-Cas9 derived *knockouts* of NUMA1 in both HAP1 cells and non-small lung cancer (NSLC) cell lines. Our results highlight a mechanism relying on mitotic arrest upon loss of BMI1 and a new genetic resistance mechanism. These observations add further knowledge to the complex and context-dependent involvement of BMI1 in cancer. Our findings contribute to a better understanding of BMI1-associated pharmacological cancer treatment strategies.

## Results

### Loss of BMI1 induces cell death in cancer cell lines

In order to test BMI1-dependent lethality in HAP1 cells, we transduced doxycycline-responsive plasmid systems into wild-type HAP1 cells to generate an inducible expression of shRNA targeting *BMI1* (shBMI1) or control shRNA (shRND). We assessed knockdown efficiency upon doxycycline treatment by comparing RNA and protein levels at different time points. As expected, doxycycline-induced shBMI1 resulted in a decreased BMI1 expression, both at RNA (avg knockdown, 93%) and protein level after 48 hours of doxycycline treatment (Fig 1A and 1B).We next evaluated HAP1 cell viability upon shBMI1 induction through colony formation, cell count, and cell viability assays. shBMI1-induction reduced HAP1 cell proliferation after doxycycline exposure, compared with shRND-induced HAP1 cells, indicating *BMI1* knockdown is lethal in HAP1 cells (Fig 1C, 1D, **and S1 Fig)**.

**Fig 1.**
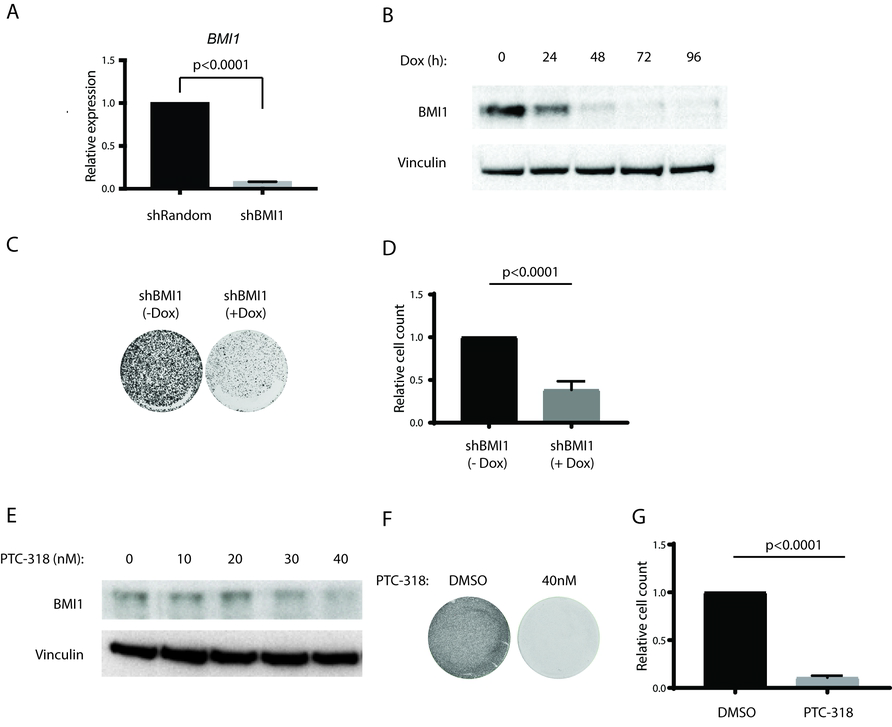
Genetic knockdown and pharmacological inhibition of BMI1 induces cell lethality. (A) Relative mRNA expression levels of BMI1 HAP1 cells transduced with inducible shRNA constructs FH1t-Random (shRandom) or FH1t-BMI1 (shBMI1) 96 hours after doxycycline treatment. (B) Protein expression levels of BMI1 in HAP1 cells transduced with shBMI1 0-96 hours after doxycycline treatment. Vinculin was used as a loading control. (C) Cell survival upon BMI1 knockdown was confirmed in HAP1 cells transduced with shBMI1 through a colony formation assay one week after plating with or without doxycycline treatment or (D) through relative cell counts 96 hours after plating with or without doxycycline. (E) Protein expression levels of HAP1 cells treated with different concentrations of PTC-318. Vinculin was used as a loading control. (F) Cell survival of HAP1 cells treated with PTC-318 through colony formation assay one week after treatment with DMSO (0.1%) or PTC-318 (40 nM) and (G) relative cell counts 48 hours after treatment with DMSO (0.1%) or PTC-318 (40 nM). Error bars represent SD. Student’s t test was performed for statistical testing.

In order to use a pharmacological setting, and hence more clinically-relevant, we inhibited BMI1 expression post-transcriptionally with the small molecule AB057609107 (PTC-318). PTC-318 was identified by performing a high-throughput compound screening through the gene expression modulation by small molecules (GEMS) technology. A luciferase open reading frame flanked by the untranslated region (UTR) of BMI1 was used as a reporter to screen for small molecules inhibiting BMI1 at the transcriptional level.

To avoid inhibitor-derived side-effects, we screened for the lowest concentrations of PTC-318, leading to cell death. PTC-318 titrations showed that treatment with low nanomolar concentrations (>20 nM) resulted in reduced BMI1 protein expression compared with dimethyl sulfoxide (DMSO) treated HAP1 control (Fig 1E). Moreover, 20nM and 40nM PTC-318 treatment-induced cell death as measured by colony formation and cell proliferation assays (Fig 1F, 1G, **and S1 Fig)**. These results suggest that similar to BMI1 knockdown, the pharmacological inhibition of *BMI1* with nanomolar concentrations of PTC-318 induces lethality in HAP1 cells.

In order to confirm that PTC-318-induced lethality was caused by BMI1 regulation and to exclude potential off-target toxicity, we expressed BMI without its UTR in HAP1 cells. These cells showed increased BMI1 protein levels compared with wild-type HAP1 cells. Importantly, upon treatment with PTC-318, cells ectopically expressing BMI1 showed increased resilience to PTC-318 treatment compared with wild-type HAP1 cells (Fig 2A and 2B), suggesting that BMI1 transcriptional inhibition is responsible for PTC-318 cytotoxicity.

**Fig 2.**
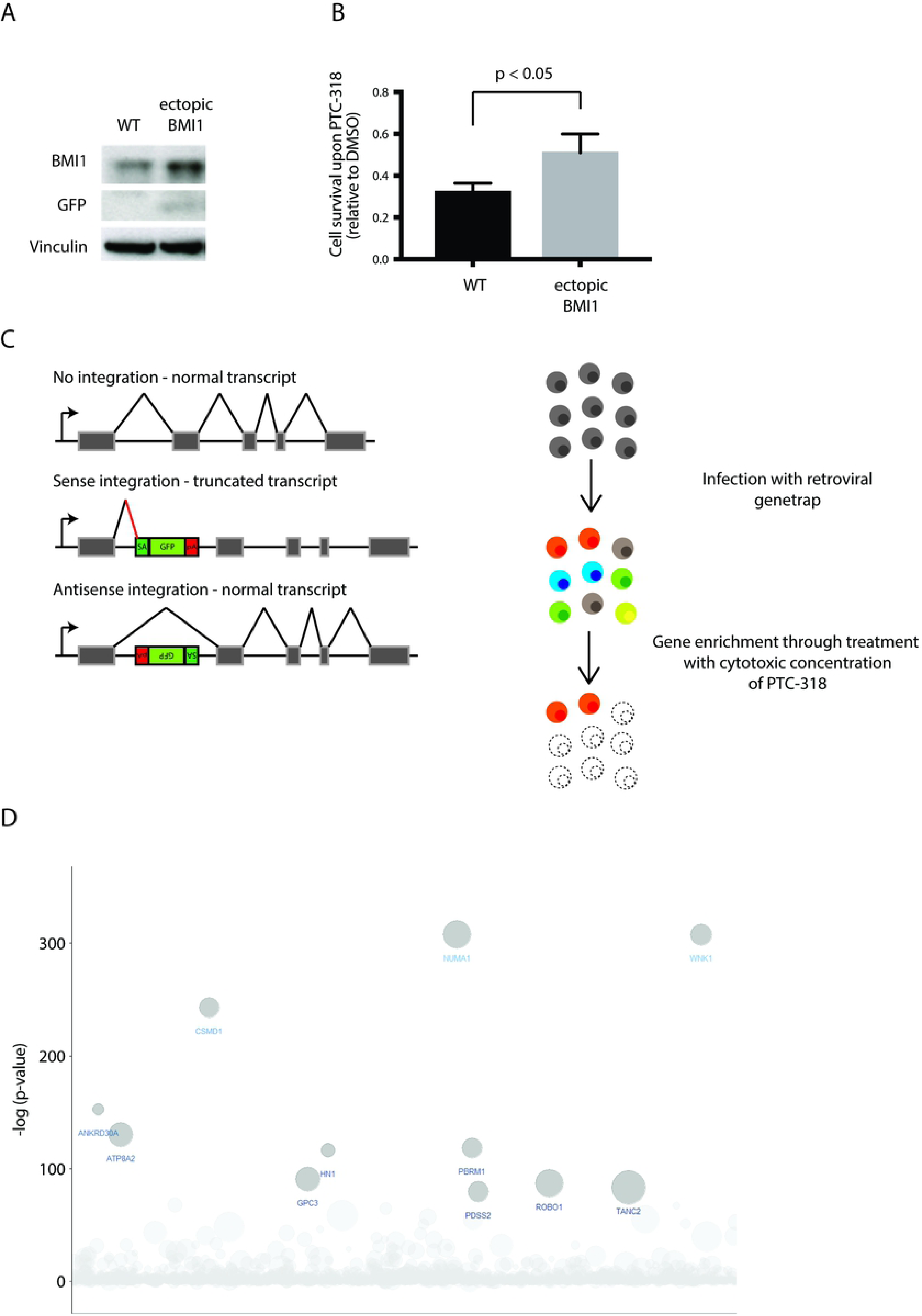
A haploid genetic screen shows enrichment of NUMA1 mutations upon BMI1 inhibition with PTC-318. (A) BMI1 and GFP protein expression of wildtype HAP1 cells and HAP1 cells ectopically expressing BMI1 in the absence of 3’UTR through the transduced FUGW-BMI1 construct. (B) HAP1 cell survival relative to corresponding DMSO-control 48 hours after treatment with 20 nM PTC-318. Error bars represent SD. Student’s t test was performed for statistical testing. (C) Schematic overview showing insertional mutagenesis of HAP1 cells using a retroviral gene trap (left) and enrichment screening process (right). (D) Bubble plot depicting genes enriched for unique gene-trap insertions in HAP1 cells treated with 40 nM PTC-318. The y-axis shows the significance and the x-axis the genes for which the gene-trap insertions were mapped in alphabetical order. Size of the bubble corresponds to the number of unique inactivating gene-trap insertions.

### NUMA1-integrations cause resistance to PTC-318-derived cell death in HAP1 screen

Given the specificity and potent cytotoxicity of PTC-318, we performed an insertional mutagenesis drug resistance screen in HAP1 cells to find new BMI1-associated interactions and resistance mechanisms. Due to their haploid genome, HAP1 cells are particularly amenable to mutagenesis as only a single allele requires inactivation to obtain a loss-of-function. This approach has been successfully used to investigate various biological processes, including virus entry (20–22), T cell-mediated killing (23), and drug response (24,25) in both haploid (HAP1) and near-haploid (KBM7) human cells.

In essence, wild-type HAP1 cells were transduced with a retroviral GFP gene trap that randomly integrates into the genome. The gene trap most commonly integrates into introns and can land in two orientations: sense or antisense. Owing to the unidirectional design of the gene trap sense-orientation integrations are likely to produce truncated and impaired transcription products. (Fig 2C).

We subjected mutagenized HAP1 cells to a stringent selection with 40 nM of PTC-318 for two weeks. Next, the genomic DNA of surviving HAP1 cells was isolated, and the insertion sites were subsequently amplified and sequenced. Gene trap integrations were then assigned to genes and counted as inactivating integrations if inserted in sense with the transcriptional orientation in an intron or regardless of orientation in an exon. To identify genes enriched in the PTC-318 treated population, we compared the number of disruptive integrations per gene in our experimental dataset with a non-treated control dataset using a one-sided Fisher’s exact test. We applied a stringent cut-off and considered all genes with an FDR-corrected p-value lower than 10^−5^ as enriched for inactivating integrations.

Interestingly, we found several reported antagonists of Polycomb repressors among the most significant hits required for cell survival, including two genes (*PBRM1* and *ARID2*) encoding essential PBAF subgroup proteins of the nucleosome remodeling complex SWI-SNF, (Fig 2D **and S1 Table)**. Strikingly, the most significantly enriched gene in the HAP1 population, *NUMA1* (p<0.0001) had not previously been associated with *BMI1*. *NUMA1* encodes a 238 kDa nuclear mitotic apparatus protein, which is mainly reported to be involved in dynein-associated tethering of spindle pole microtubules during mitosis, recruiting dynein to the cell cortex (26–28). NUMA1 has also been implicated in a conserved mechanism establishing asymmetric cell division associated with differentiation. The ` features associated with NUMA1, as well as its tumor suppressive and cell differentiating properties (29–32), suggested it as an attractive BMI1-antagonizing candidate, especially considering the role of BMI1 in maintaining cell stemness in healthy and cancer cells. To understand this new genetic association, we decided to focus on *NUMA1*-loss as a possible resistance-inducing mechanism against BMI1 inhibition.

### Knockout of *NUMA1* in HAP1 cells induces resistance to cell death upon BMI1 inhibition

As validation of the screen results, we checked whether loss of *NUMA1* expression would result in resistance to the lethality induced by BMI1 inhibition. We generated clonal *NUMA1* knockout HAP1 cell lines (NUMA1-KO) by targeting the first exon of *NUMA1* with CRISPR-Cas9. (Fig 3A and 3D). We found that NUMA1-KO cells were significantly more resistant to the toxicity of shRNA knockdown of BMI1 or PTC-318 treatment compared with wildtype HAP1 (hereafter named NUMA1-WT) cells (Fig 3B and 3E). We found a more than 2-fold higher cell viability in NUMA1-KO cells compared to WT upon BMI1 knockdown and inhibition, respectively, following quantification of Alamar blue-stained cells (Fig 3C and 3F).

**Fig 3.**
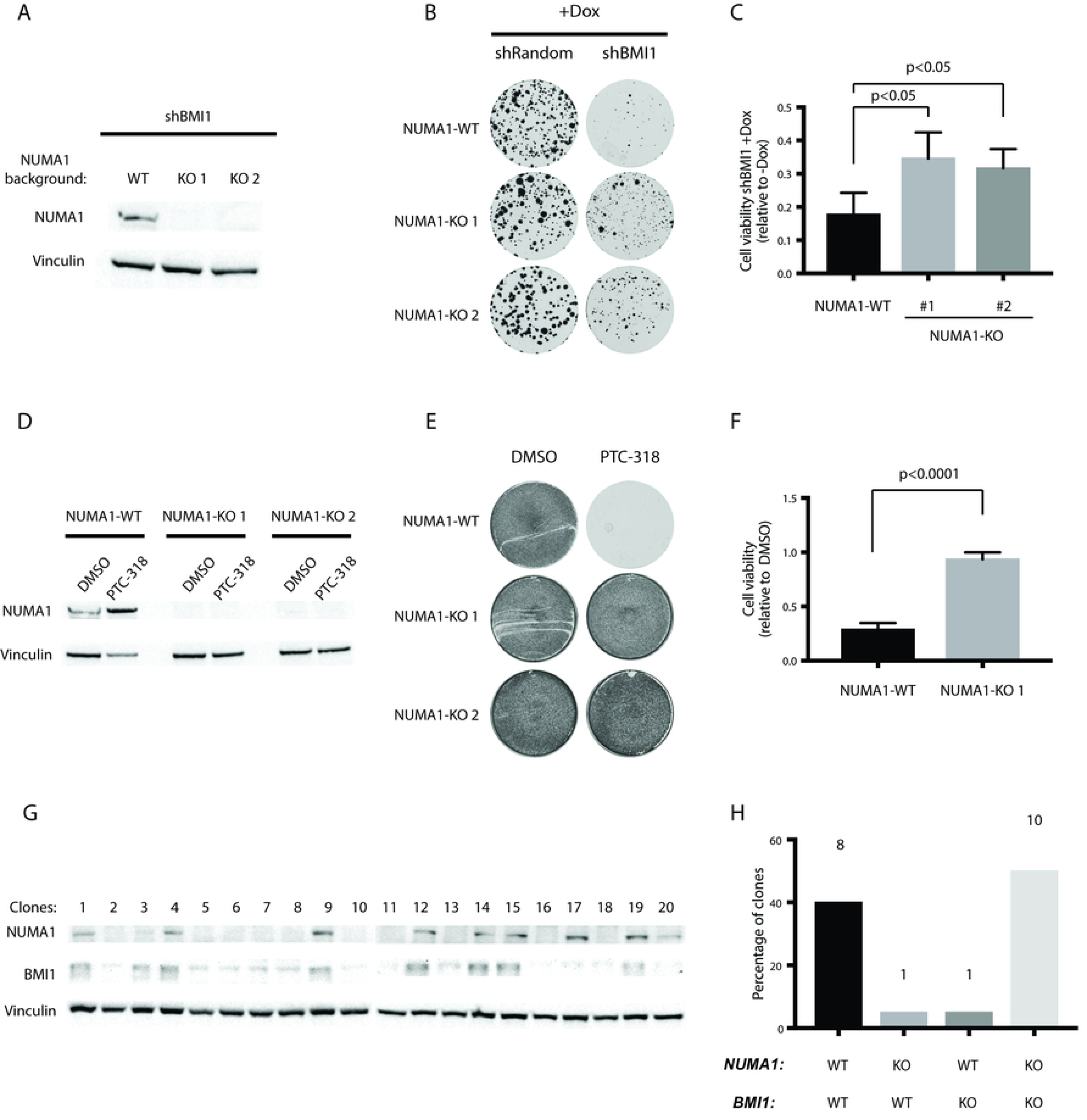
Cas9-induced NUMA1 knockout rescues HAP1 cytotoxicity caused by BMI1 inhibition. (A) Protein expression of NUMA1 in populations of HAP1 cells transduced with a doxycycline inducible shBMI1 construct in NUMA1-WT or NUMA1-KO backgrounds. (B and C) Survival of the HAP1 NUMA1-WT and NUMA1-KO clones transduced with the inducible shBMI1 construct and maintained with or without doxycycline (B) colony formation and (C) cell viability. (D) Expression of NUMA1 protein in HAP1 NUMA1-WT and NUMA1-KO clones treated either with 0.1% DMSO or 40 nM of PTC-318. The survival of these cells was analyzed by (E) colony formation assay and (F) cell viability. (G) Protein expression of NUMA1 and BMI1 in NUMA1-WT clones, transfected with CRISPR-Cas9 targeting BMI1, single-cell sorted, and expanded (left). (H) Percentages of each genetic background based on the respective protein expression (right). Error bars represent SD. Student’s t test was performed for statistical testing.

In order to further validate that the NUMA1-KO resistance was specific to BMI1 inhibition, we attempted to establish BMI1 knockout (BMI1-KO) HAP1 clones. We transfected NUMA1-WT or NUMA1-KO HAP1 cells with CRISPR-Cas9 targeting *BMI1*, hypothesizing that NUMA1-WT cells would survive the BMI1-KO to a lesser extent compared with NUMA1-KO cells. While we were able to establish ten BMI1-KO clones in NUMA1-KO background, we retrieved only one NUMA1-WT clone with reduced BMI1 expression (Fig 3G and 3H).

These results indicated that loss of NUMA1 results in resistance to the toxicity resulting from loss of BMI1, whether induced by shRNA, inhibitor, or CRISPR-Cas9. It also further confirmed the BMI1-specific effect of PTC-318 and that the NUMA1-derived resistance is specific to BMI1-associated lethality.

### BMI1-loss induces mitotic arrest and cell death

Since NUMA1 is important for proper mitotic progression, we assessed the cell cycle profiles of HAP1 cells treated with a lower concentration of PTC-318 (20nM). We chose the lower concentration in order to maintain target engagement and limit loss of cell viability.

Both genetic knockdown and pharmacological inhibition of BMI1 resulted in increased numbers of cells arrested in mitosis, as shown by the increase in Histone H3 phosphorylation (pH3). Compared with shRND or DMSO controls, BMI1 knockdown and PTC-318 treatment resulted in a 2-fold and 8-fold increase in pH3 positive cell populations, respectively (Fig 4A).

**Fig 4.**
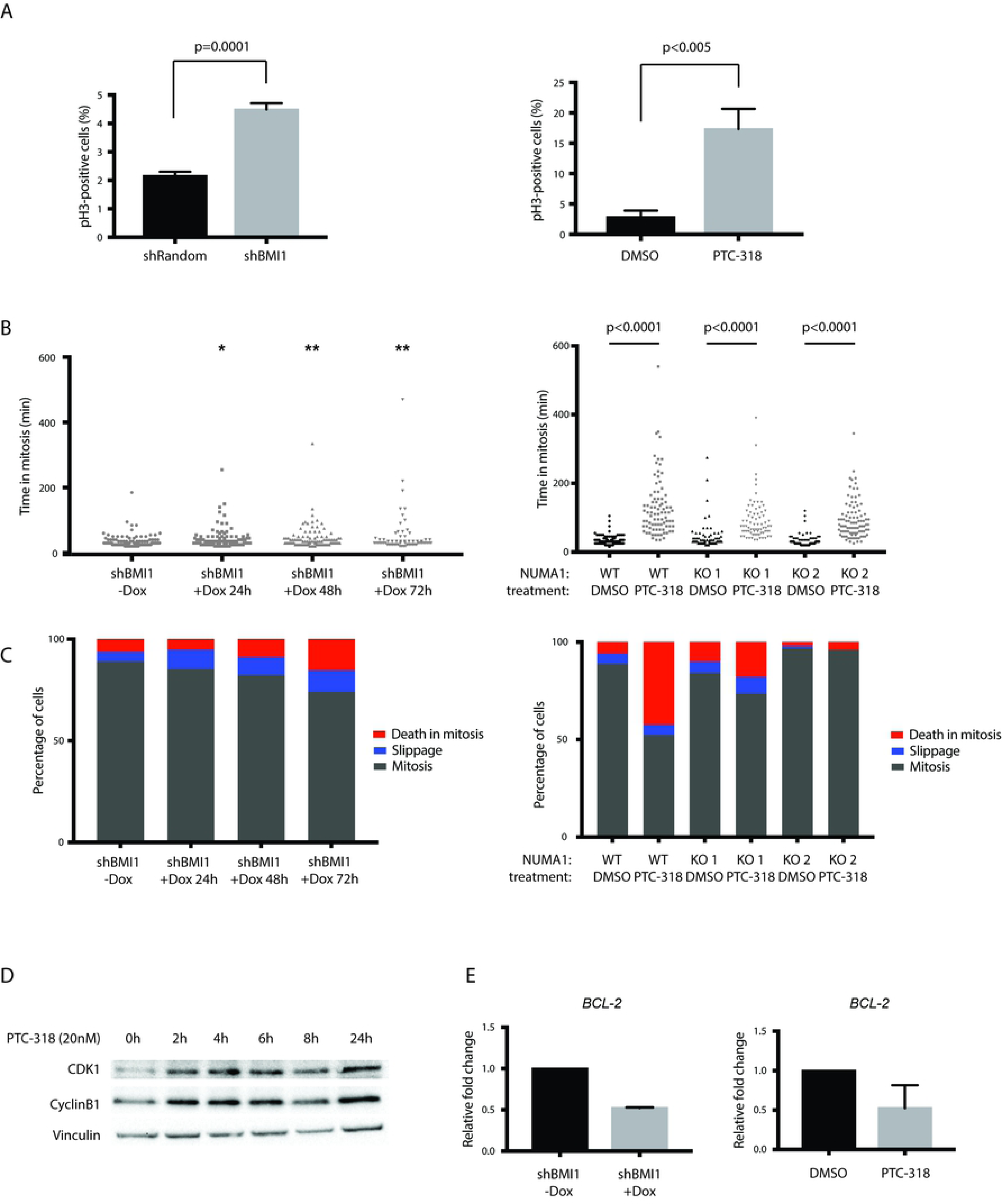
BMI1 inhibition induces mitotic cell death in NUMA1-WT cells but not in NUMA1-KO. (A) Quantification of phospho-histone H3 populations of HAP1 cells transduced with the inducible shRandom or shBMI1 constructs where cells were treated with doxycycline for 96 hours before measurement (left graph), and cells treated for 48 h with 0.1% DMSO or 40 nM PTC-318 (right graph). (B) Scatter dot plot representation of time in mitosis (from nuclear envelope breakdown to anaphase onset) of HAP1 cells transduced with the inducible shBMI1 construct and treated with doxycycline for 0-96 hours (left graph), or NUMA1-WT and NUMA1-KO cells treated with 20 nM PTC-318 (right). Bars represent the mean and standard deviation (SD). (C) Analysis of the mitotic fate of cells analyzed in B). The graphs depict the percentage of cells in mitosis (dark grey), mitotic slippage (blue), and death in mitosis (red) (D) Protein expression of CDK1 and CyclinB1 in HAP1 cells at different time points after treatment with 40 nM of PTC-318. (E) Relative expression levels of BCL-2 in HAP1 transduced with inducible shBMI1 construct (with or without doxycycline for 96 hours; left) or treated with 0.1% DMSO or 20 nM PTC-318 for 4 hours (right). Error bars represent SD. Student’s t test was performed for statistical testing (*p<0.05 and **p<0.01).

Having confirmed mitotic arrest induced by BMI1 inhibition, we next followed the mitotic progression of HAP1 by live-cell imaging upon doxycycline treatment at different time-points (24, 48, and 72 hours) or 20 nM PTC-318 treatment. We found a modest albeit significantly increased time spent in mitosis in all doxycycline-induced shBMI1 HAP1 cells (41, 45, and 53 minutes, respectively) compared with control cells (34 minutes) (Fig 4B **and S2 Fig)**.

In contrast to the modest increases of time spent in mitosis observed in shBMI1 cells, analysis of NUMA1-WT HAP1 cells treated with PTC-318 showed a more prolonged mitotic arrest compared with DMSO-treated cells. While the mitotic duration of DMSO-treated cells was 31 minutes, PTC-318 treated NUMA1-WT cells spent 131 minutes in mitosis. In general, we observed that BMI1 inhibition with PTC-318 resulted in a robust mitotic arrest in both NUMA1-WT and NUMA1-KO cell lines. However, the arrest was longer in NUMA1-WT than in NUMA1-KO cell lines (131 minutes and 92 minutes, respectively).

After observing mitotic arrest induced by BMI1 inhibition, through both shBMI1 and PTC-318, we examined the fate of these cells by live-cell imaging (Fig 4C). While most of the untreated cells underwent chromosome segregation and completed cell division, we observed an enrichment of mitotic cell death upon BMI1 inhibition. This effect was more pronounced in PTC-318-treated cells than in the shBMI1 population.

It has been described that the balance between two competing networks dictates cell fate. While the CyclinB1/cyclin-dependent kinase 1 (CDK1) complex is essential for mitotic progression, opposing signals, regulating, for example, the apoptosis-regulating protein B-cell lymphoma 2 (BCL-2), can induce mitotic cell death. The fate of a cell in mitotic arrest is thus linked to the antagonizing levels and timing of these two independent networks: if cell death signals increase, CyclinB1 levels decrease (33).

Thus, we measured the expression levels of CyclinB1/CDK1 and BCL-2 in BMI1-inhibited HAP1 cells. Consistent with the observed mitotic arrest, both CDK1 and CyclinB1 protein levels were elevated at early time-points after PTC-318 treatment (between 2 and 4 hours), and remained high up to 24 hours after treatment (Fig 4D). In agreement with these results, live-cell imaging data showed that the onset of mitotic arrest was around 3-4 hours after treatment **(S3 Fig)**. In order to quantify cell death signals induced by BMI1 inhibition, we determined RNA levels of the anti-apoptotic marker *BCL-2* and observed a significant decrease after PTC-318 treatment compared with DMSO-treated control cells (Fig 4E). Similar to PTC-318 treatment, induction of shBMI1 in the HAP1 cells significantly decreased *BCL2* expression. Together, these results suggest that BMI1 inhibition induces mitotic arrest in HAP1 cells, ultimately causing cell death in the NUMA1-WT background but not in the NUMA1-KO population.

### Inhibition of BMI1 induces cell death in NSCLC cell lines in mitosis

We asked whether toxicity associated with BMI1-inhibition was could be relevant in other cancer cell types and rule out that our findings were limited to haploid cells. Previous studies have demonstrated that BMI1 is overexpressed in non-small cell lung cancer (NSCLC) and that its expression was associated with NSCLC progression (34–36). We, therefore, tested whether NSCLC would be sensitive to BMI1 inhibition. We treated NSCLC cell lines h520 and SK-MES-1 with PTC-318 and found that, as in HAP1 cells, low nanomolar concentrations of PTC-318 resulted in a significant reduction in cell survival of both lines (IC50 h520: 1.9 nM) (IC50 SK-MES-1: 7.8 nM) compared with untreated controls. (Fig 5A and 5B). Both NSCLC cell lines showed a significant accumulation pH3 mitotic marker upon PTC-318 treatment, further suggesting that our findings in the HAP1 population can be generalized to other cancer types (Fig 5C and 5D).

**Fig 5.**
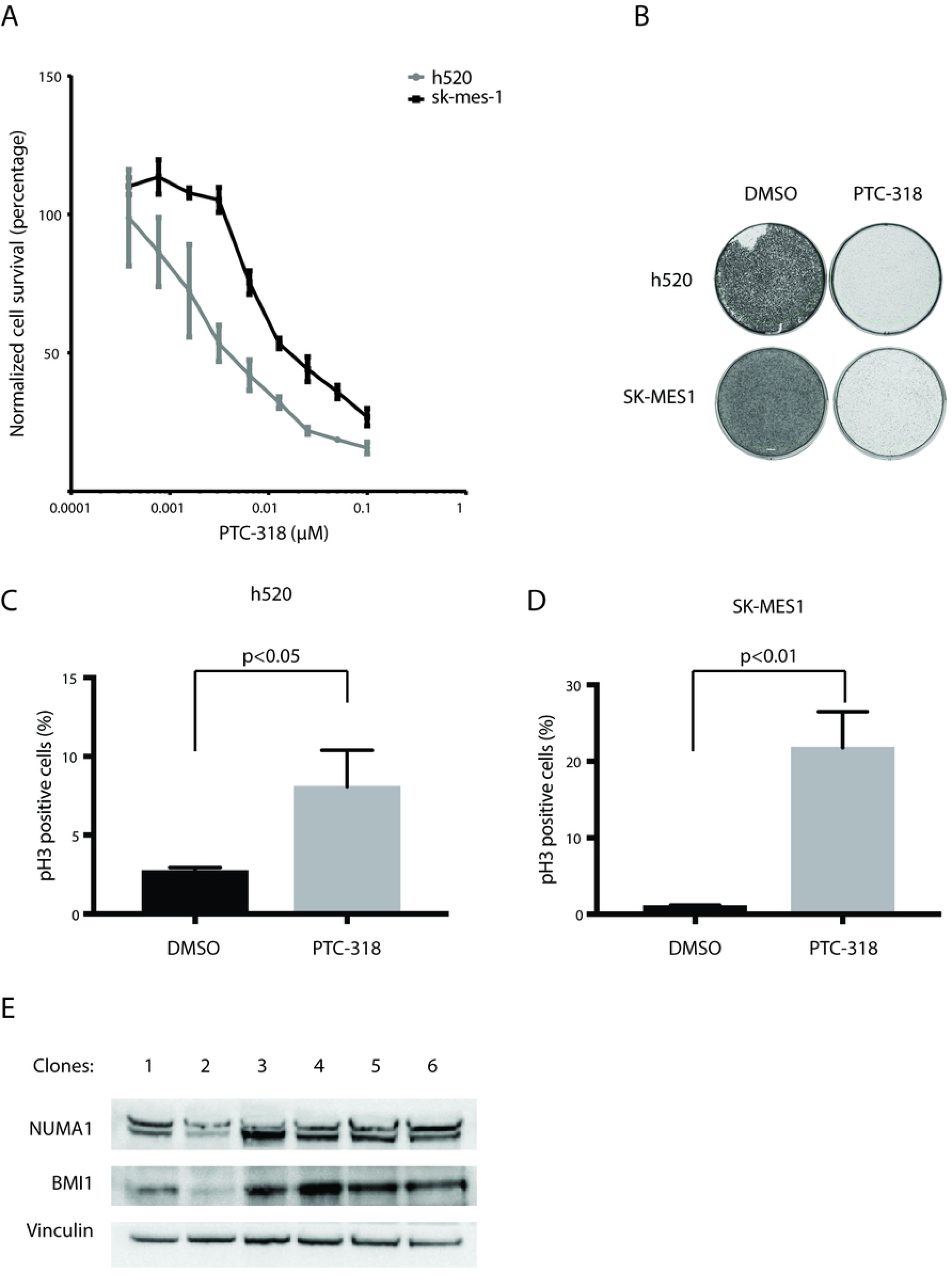
Non-small cell lung cancer cell lines are sensitive to BMI1 inhibition. (A) IC50 measurements of NSCLC cell lines h520, SK-MES-1, and SWI-1573. X axis shows the PTC-318 concentration in uM and y axis the percentage of surviving cells relative to non-treated control counterparts. (B) Colony formation of h520 and SK-MES-1 cells treated with either 0.1% DMSO or 40 nM of PTC-318. (C) Percentages of h520 phospho-histone H3 populations after treatment with either 0.1% DMSO or 40 nM PTC-318 determined by flow cytometry quantification. (D) Percentages of h520 phospho-histone H3 populations after treatment with either 0.1% DMSO or 80 nM PTC-318 determined by flow cytometry quantification (E) Protein expression of NUMA1 and BMI1 in NUMA1-WT clones of a h520 population transfected with Cas9 targeting BMI1, single-cell sorted, and expanded. Error bars represent SD. Student’s t test was performed for statistical testing.

Finally, we validated the essential function of BMI1 in NSCLC as we did for HAP1 cells. We transfected the two NSCLC cell lines with CRISPR-Cas9 targeting *BMI1* and sorted single-cell clones. While we were not able to grow any single clones from the transfected SK-MES-1 population, we retrieved six h520 clones after single-cell sorting. Five of these clones showed regular expression of BMI1, and one showed reduced BMI1 expression (Fig 5F). Interestingly, the clone with the lowest BMI1 expression also showed reduced levels of NUMA1 compared with the other h520 cell clones. These observations are in line with our main finding that BMI1 inhibition is tolerated only in the context of NUMA1 knockout or at least NUMA1 reduction, while it is lethal in NUMA1-proficient cells.

Altogether, we were able to show mitotic arrest and cytotoxicity caused by BMI1 loss in two commonly used NSCLC in vitro models, indicating that the observed phenotypes related to BMI1 inhibition extend to non-haploid cancer cells as well.

## Discussion

Loss of BMI1 has been shown to induce growth arrest and cell death in cancer cells both *in vitro* and *in vivo* (37–39). However, although BMI1 inhibition leads to impaired cell proliferation in different systems, the exact mechanisms are often context-dependent. Also, while BMI1 expression can cause resistance against different cancer treatment strategies, including chemotherapy or radiation (15–18), little is known about possible resistance-inducing responses to cell death resulting from BMI1 inhibition.

Encouraged by the therapeutic potential of BMI1 inhibition, we assessed the effect of PTC-318, a new and previously unpublished small molecule inhibitor of BMI1, on human cancer cells and evaluated its underlying mechanisms of action. We found that chemical inhibition of BMI1 was very efficiently causing cell death in human haploid HAP1 cells – a cell line originally derived from chronic myeloid leukemia cells (40) – and in non-small cell lung cancer (NSCLC) cell lines.

A PTC-318 resistance screen on HAP1 cells allowed us to address genetic alterations underlying resistance to BMI1 inhibition, thus expanding our understanding of previously unknown BMI1 mechanisms. Reassuringly, our screen presented significant hits including genes of the SWI-SNF complex, one of the most studied ATP-dependent chromatin remodeling complexes, known to regulate and antagonize the function of polycomb repressive complexes (41). Among these hits, we found PBAF-specific genes PBRM1 (BAF180) and ARID2 (BAF200) from the Polycomb-antagonistic SWI-SNF complex. While the finding of the two independent components of the PBAF chromatin remodeling complex might be expected based on the known dose-dependent antagonistic relationship between Polycomb repressors and SWI/SNF remodelers (41), Nuclear Mitotic Apparatus proteins (*NUMA1*) was an unexpected candidate as the most significant hit in our HAP1 screen. Here, we confirmed that loss of NUMA1 established by CRISPR-Cas9 could rescue the lethality induced by shRNA-established BMI1 knockdown and PTC-318 inhibition.

Using different genetic approaches, including genetic interference through shRNAs and ectopic BMI1 expression, we could confirm that PTC-318-derived cell death was on-target and caused by BMI1 inhibition. It is worth noticing that knockdown of *BMI1* by shRNA resulted in significant, but weaker phenotypic changes in our studied cell lines compared with PTC-318 treatment. The relatively weaker phenotypes might be because the shRNAs only achieved partial inhibition of BMI1 in comparison with PTC-318 treatment. We cannot, however, entirely exclude that PTC-318 may have some additional effects beyond BMI1 inhibition.

Apart from the similarities observed between shRNA and small molecule-induced BMI1 inhibition, we also demonstrated that our attempts to establish BMI1 knockout lines were unsuccessful in NUMA1-WT backgrounds, but not in NUMA1-KO backgrounds confirming BMI1-dependency in multiple cell lines. Interestingly, the only clone surviving BMI1 knockout in the NSCLC cell line h520 expressed lower levels of NUMA1, further underscoring the genetic interactions between NUMA1 and BMI1.

NUMA1 and BMI1 are involved in several shared cellular processes that could suggest possible direct or indirect mechanistic relationships between the two proteins. For example, just as BMI1, NUMA1 is involved in the DNA damage repair and homologous recombination (42), cell differentiation (30–32), higher-order chromatin organization (43), and cell cycle progression. The cell cycle properties of NUMA1, in particular in mitosis, were especially interesting to us since previous studies have implicated the EZH2 protein of polycomb repressive complex 2 (PRC2) as a regulator of cell cycle checkpoints, including cyclin-dependent kinase inhibitor p21 (44). According to the findings, loss of EZH2 in cancer cells abrogated cell cycle arrest in G1 and G2/M upon treatment with DNA-damaging agents Adriamycin or etoposide. Importantly, several studies have reported the involvement of BMI1 in mitosis or G2/M phase (45,46). Wei et al. analyzed the DNA damage repair mechanisms in breast cancer MCF7 cells upon ectopic expression or inhibition of BMI1. They found that while ectopic BMI1 expression resulted in reduced G2/M arrest upon etoposide-treatment, knockdown of BMI1 had the opposite effect.

In light of our screening results and the reported role of NUMA1 in the cell cycle, we further validated that inhibition of BMI1 in *NUMA1* proficient cells resulted in a mitotic arrest followed by cell death in mitosis, a phenotype that could be rescued by inactivation of *NUMA1*. The mitotic arrest observed in our study upon inhibition of BMI1 coincided with increases in CDK1 and Cyclin-B1 protein levels. The findings further support previous studies demonstrating a CDK1-induced switch from mitotic arrest to apoptosis (47). According to their results, this apoptotic switch is derived from the phosphorylation of Bcl-2/Bax family proteins upon interfering microtubule formation. In our study, both genetic and drug-induced BMI1 regulation decreased RNA expression of *BCL-2*, a known protein with anti-apoptotic functions (48), suggesting deregulation of both mitosis and cancer-associated survival mechanisms upon BMI1 inhibition. Still, our results from the cell cycle profiling indicate that knockout of NUMA1, while rescuing the lethal phenotype induced by BMI1 inhibition, does not rescue mitotic arrest. The latter observation may suggest that BMI1 directly or indirectly acts as a checkpoint during improper mitotic progress caused by an impaired *NUMA1* expression.

In conclusion, our results investigating the effects of PTC-318 on cancer cell lines indicate that inhibition of the oncogene BMI1 induces cancer cell death, making it a potential approach to treat certain types of cancers – either as a single agent or in combination with other inhibitors. The emergence and testing of new BMI1 inhibitors for the treatment of cancer suggest that BMI1 is a relevant target for cancer therapy. In our study, we unveiled how the inhibition of BMI1 induces cell death through mitotic arrest that was successfully rescued by the depletion of NUMA1 expression. In our experimental set-up, *NUMA1*-mutations strongly associated with cancer cell survival and may have an essential role – and hence serve as a potential biomarker – in resistance mechanisms in treatments associated with BMI1 inhibition. Although further studies are needed to establish a complete picture of the pathways and mechanisms linked with BMI1 overexpression and inhibition in cancer, our findings highlight an essential and novel mechanism of BMI1 in cancer cells, which could contribute to the development of effective cancer therapies.

## Materials and Methods

### Cell culture

HAP1 cells have been described previously (49). They and were maintained in IMDM + GlutaMAX (Gibco) supplemented with 10% fetal calf serum (FCS; Sigma-Aldrich), and penicillin–streptomycin (Gibco). NSCLC cell lines h520 and SK-MES-1 (provided by the A. Berns laboratory at the Netherlands Cancer Institute, Amsterdam, the Netherlands) were maintained in DMEM/F12 + GlutaMAX (Gibco) supplemented with 10% FCS and penicillin–streptomycin (Gibco).

### Haploid genetic screen

Procedures for the generation of gene-trap retrovirus and HAP1 mutagenesis have been described previously (22,49). To select HAP1 variants resistant to BMI1 inhibitor PTC-318, approximatively 10^8^ mutagenized HAP1 cells (>90% haploid) were seeded in fourteen T175 cell culture flasks. The cells were exposed to 40 nM of PTC-318 24 hours after seeding and incubated for fourteen days. Surviving HAP1 clones were trypsinized and washed before amplification and analysis of integration sites, as described in (22,50). In brief, as a first step insertion sites were amplified in a linear amplification reaction using a biotinylated primer. Products were captured on streptavidin-coated magnetic beads, washed, and subjected to single-stranded DNA linker ligation followed by a second PCR to finalize the products for Illumina sequencing. Subsequently, as described for enrichment mutagenesis screens in Staring et al., 2017, sequence reads were aligned to the human genome (HG19) and uniquely aligning insertion sites were mapped to the genomic coordinates (RefSeq) of non-overlapping protein-encoding gene regions. Gene trap integrations in sense in introns, or regardless of orientation in exons, were considered disruptive. To select genes enriched for mutations after PTC-318 selection, the number of disruptive integrations in each gene was compared to those retrieved in an unselected HAP1 population (22) using a one-sided Fisher’s exact test and corrected for multiple testing (Benjamini and Hochberg FDR).

### Generation of knockdown, overexpression, and knockout cell lines

For BMI1 knockdown experiments, we used doxycycline-inducible FH1-tUTG-RNAi vectors (51), as described previously (52). For ectopic expression of BMI1, we used the BMI1-overexpression vector with FUGW vector backbone (FUGW-BMI1; Addgene 21577) Cells in 10-cm plates were transduced using 10 μg of FH1-tUTG-RNAi vector or FUGW-BMI1, 3.5 μg VSV-G, 2.5 μg REV, and 5 μg pRRE in CaCl_2_ (2.5M). GFP positive cells were sorted by flow cytometry (MoFlo), collected in respective media containing 20% FCS, and subsequently spun down and plated in media containing 10% FCS. shRNA targeting sequences: shBMI1, 5′-GGAGGAGGTGAAGTATAAA′; shRandom, 5′-ATTCTTACGAAACCCTTAG-3′.

For Cas9-induced knockout, we used SpCas9 and chimeric guide RNA (gRNA) expression plasmid pX330-U6-Chimeric_BB-CBh-hSpCas9 (Addgene 42230) encoding gRNAs targeting NUMA1 (NuMA-KO) or BMI1 (BMI1-KO). Cells were transfected in 6-well plates with 1.2-1.6 μg of Cas9-gRNA construct together with 10% 1.5 µg of mPB-L3-ERT2.TatRRR-mCherry plasmid, following the Lipofectamine 2000 protocol. mCherry-positive cells were single-cell sorted by flow cytometry (MoFlo) into 96-well plates 48 hours after transfection and incubated for two weeks before expansion. Gene mutations were validated by Sanger sequencing and western blot analysis. gRNA targeting sequences were 5′-GACACTCCACGCCACCCGGG-3′ for NuMA-KO and 5′-AACGTGTATTGTTCGTTACC-3′ for BMI1-KO.

### Western blot analysis

Whole-cell extracts pelleted and prepared in RIPA buffer (50 mM Tris, pH 8.0, 50 mM NaCl, 1.0% NP-40, 0.5% sodium deoxycholate, and 0.1% SDS) containing protease inhibitor cocktail (Complete; Roche) and phosphate inhibitors (10 mM Na fluoride final concentration, 1 mM sodium orthovanadate final concentration, and 1 mM NaPPi final concentration). Equal amounts of protein, as determined by a Bio-Rad Protein Assay Dye Reagent on Nanodrop 2000c, were resolved on NuPage-Novex 4–12% Bis-Tris gels (Invitrogen) and transferred onto nitrocellulose membranes (0.2 m; Whatman). Membranes were blocked in phosphate-buffered saline (PBS) with 0.1% Tween-20 (PBST) and 5% BSA for 1 h, incubated with primary antibodies in PBST 1% BSA overnight at 4°C, and incubated with secondary antibodies coupled to HRP for 45 min in PBST 1% BSA at room temperature. Membranes were imaged on a BioRad ChemiDoc XRS+. The following antibodies were used for western blot analyses: anti-BMI1 D20B7 (Cell Signalling) anti-GFP (Abcam, ab6556), anti-NuMA (Thermo Fisher, PA5-22285), anti-CDK1 (Bethyl, A303-664A), Cyclin B1 (GNS1; Santa Cruz, sc-245).

### Gene expression

Total RNA was isolated with the ReliaPrep™ RNA Cell Miniprep System (Promega) according to the manufacturer’s instructions. RNA quantity and quality were assessed using a Nanodrop 2000c (Thermo Scientific). Primers details are available upon request.

### Clonogenic assays

To test PTC-318 lethality, 400,000 HAP1 cells/well or 100,000 NSCLC cells/well were seeded in 2 ml/well in 6-well plates and incubated over night at 37°C. Dimethyl sulfoxide (DMSO) diluted PTC-318 (80 μM) was further diluted with the corresponding medium to obtain the desired concentrations. For control treated cells, DMSO was diluted to match the DMSO-concentrations of PTC-318 dilutions (below 0.1%). PTC-318 or DMSO dilutions were distributed to the cells (2 ml/well) and incubated for one week at 37°C. Cells were next washed with PBS and incubated with 0.5 ml of 0.1% crystal violet (50% methanol) for 10 min at room temperature. Imaging of the wells was performed with GelCount (Oxford Optronix).

FH1t-BMI1-induced lethality was assessed by plating 100,000 HAP1 cells/well in 6-well plates in cell medium with or without doxycycline. Doxycycline treatment was performed every day until the end of the experiment (96 hours). For time-point experiments, all evaluated time-points were plated at 0 hours and assessed 96 hours after plating to maintain similar confluency. Doxycycline treatment start depended on conditions tested (96 h doxycycline: 0 h after seeding, 72 h: 24 hours after seeding, 48 h: 48 hours after seeding, and 24 h: 72 hours after seeding). Cells were washed with phosphate-buffered saline (PBS) and incubated with 0.5 ml crystal violet for 10 min at room temperature. Imaging of the wells was performed with a GelCount.

### Cell viability assay

To assess cell viability upon PTC-318 treatment, 40,000 HAP1 cells/well or 20,000 HAP1 cells/well were plated in 24-well plates (500 μl/well) and incubated for 24 hours at 37°C. Next, the cells were treated with respective concentrations of PTC-318 (as described above) and incubated for 48 hours. Alamar blue (10X; Invitrogen) was diluted 1:10 in corresponding cell medium and distributed to cells. Negative control cells received 3 μl of 20% sodium dodecyl sulfate (SDS). Cell viability was determined by measuring the relative fluorescence 4-6 hours after addition of alamar blue using the HP D300 Digital Dispenser.

To assess viability upon BMI1-KD, 2,000 cells were plated in 24-well plates with or without doxycycline (500 μl/well) and incubated for 24 hours at 37°C. The doxycycline was refreshed every day for one week before adding the alamar blue as previously described.

### Flow cytometry analysis

For PTC-318, 200,000 cells were plated, treated with PTC-318, and incubated for 48 hours before collection. For shBMI1, 100,000 cells were plated and incubated for 96 hours. Doxycycline was refreshed every day. After corresponding incubation times, equal amounts of cells (500,000-2,000,000) were collected after trypsinization, and thereafter centrifuged, fixed with 70% ethanol, and stored at −20°C until staining. For staining, cells were centrifuged for 3 minutes at 1,500 rpm and then washed in PBS. Cells were centrifuged again at 1,500 for 3 minutes and pellets were resuspended in 0.25% Triton X-100 (Sigma), transferred to 1.5 ml tubes, and incubated on ice for 15 minutes for permeabilization. Cells were centrifuged at 5,000 g for three minutes, resuspended in PBS containing 1% BSA and 1:100 mouse anti phospho-Histone H3 (Cell signaling, 9701), and incubated for 1 hour at room temperature. After washing, cells were resuspended in PBS containing 1% BSA and 1:300 Alexa 647 conjugated donkey anti-mouse (Thermofisher, A21238) and incubated for 30 min at room temperature and in the dark. Cells were thereafter washed, resuspended in 150 ul PBS containing 10 ug/ml DNase-free RNase A, and incubated for 30 min at 37°C. Prepared cells were stored in −20°C or 4°C until analysis on flow cytometer (LSR II). Propidium iodide (1.5 ug) was added to the samples right before assessment.

### Live cell imaging

Cells were plated on LabTek dishes and two hours before imaging, the media was changed to Leibovitz L15 CO2‐independent cell culture medium (Gibco). Mitotic progression was followed by adding SiR-DNA (Spirochrome) and drugs to the cells, that were imaged as previously described (53). In the case of shRNA-mediated knockdown of BMI1 (shBMI1), doxycycline was added to the cells 24, 48, or 72 hours prior to SiR-DNA addition. Cells were imaged every 15 minutes in a heated chamber of 37 °C using a 20x NA 0.95 air objective on an IX71 microscope (Olympus), controlled by SoftWoRx 6.0 software (Applied Precision).

## Supporting information

**S1 Fig. Inhibition of BMI1 results in reduced HAP1 cell viability.** (A) Cell survival upon BMI1 knockdown was confirmed in HAP1 cells transduced with shBMI1 through a colony formation assay one week after plating HAP1 cells transduced with either shRandom or shBMI1 and treated with doxycycline or (B) through relative cell counts of HAP1 cells transduced with shBMI1 treated with doxycycline for 72, and 96 hours after plating or without doxycycline (-Dox). (C) Cell survival of HAP1 cells treated with PTC-318 through colony formation assay one week after treatment with DMSO (0.1%) or PTC-318 (20 or 40 nM) and (D) relative cell counts 48 hours after treatment with DMSO (0.1%) or PTC-318 (20 or 40 nM). Error bars represent SD. Student’s t test was performed for statistical testing.

**S1 Table. List of the 100 most significant enriched genes after HAP1 screen exposing the cells with 40 nM PTC-318.**

**S2 Fig. Representative examples of time-lapse imaging of HAP1 cells labelled with SiR-DNA.** (A) HAP1 cells undergoing a normal mitosis with chromosome segregation (highlighted by the arrowheads). B) HAP1 cell dying after a prolonged mitotic arrest. Arrowheads point to the dying cell. C) HAP1 cell arrested in mitosis followed by slippage to interphase without DNA segregation (arrowhead). Time 0 min corresponds to nuclear envelope breakdown.

**S3 Fig. Live-cell imaging of HAP1 clones upon BMI1 inhibition.** Quantification of live-cell imaging data showing the times of individual cells. Upper three rows show cells treated with either DMSO (0.1%) or PTC-318 (20 nM) while the lower row shows HAP1 cells transduced with shBMI1 untreated (-Dox) or treated (+Dox) with doxycycline. The respective y-axes depict the individual clones.

## Acknowledgments

We thank the NKI Genomics Core Facility and the NKI Flow Cytometry Facility for their technical support. We also want to thank Vincent Blomen, Thijn R. Brummelkamp, Lorenzo Bombardelli, Anke Sparmann, and Bas Tolhuis for the valuable help, discussions, and/or constructive criticism of the manuscript.

## Author contributions

Conceptualization: Santiago Gisler and Maarten van Lohuizen.

Data curation: Gayathri Chandrasekaran.

Funding acquisition: Maarten van Lohuizen.

Investigation: Santiago Gisler and Ana Rita R. Maia.

Methodology: Santiago Gisler

Project administration: Maarten van Lohuizen.

Supervision: Maarten van Lohuizen.

Validation: Santiago Gisler and Ana Rita R. Maia.

Visualization: Santiago Gisler, Ana Rita R. Maia, and Gayathri Chandrasekaran.

Writing – original draft preparation: Santiago Gisler.

Writing – review & editing: Maarten van Lohuizen, Ana Rita R. Maia, and Gayathri Chandrasekaran.

